# An intranasal lentiviral booster broadens immune recognition of SARS-CoV-2 variants and reinforces the waning mRNA vaccine-induced immunity that it targets to lung mucosa

**DOI:** 10.1101/2022.01.30.478159

**Authors:** Benjamin Vesin, Jodie Lopez, Amandine Noirat, Pierre Authié, Ingrid Fert, Fabien Le Chevalier, Fanny Moncoq, Kirill Nemirov, Catherine Blanc, Cyril Planchais, Hugo Mouquet, Françoise Guinet, David Hardy, Christiane Gerke, François Anna, Maryline Bourgine, Laleh Majiessi, Pierre Charneau

## Abstract

As the COVID-19 pandemic continues and new SARS-CoV-2 variants of concern emerge, the adaptive immunity initially induced by the first-generation COVID-19 vaccines wains and needs to be strengthened and broadened in specificity. Vaccination by the nasal route induces mucosal humoral and cellular immunity at the entry point of SARS-CoV-2 into the host organism and has been shown to be the most effective for reducing viral transmission. The lentiviral vaccination vector (LV) is particularly suitable for this route of immunization because it is non-cytopathic, non-replicative and scarcely inflammatory. Here, to set up an optimized cross-protective intranasal booster against COVID-19, we generated an LV encoding stabilized Spike of SARS-CoV-2 Beta variant (LV::S_Beta-2P_). mRNA vaccine–primed and -boosted mice, with waning primary humoral immunity at 4 months post-vaccination, were boosted intranasally with LV::S_Beta-2P_. Strong boost effect was detected on cross-sero-neutralizing activity and systemic T-cell immunity. In addition, mucosal anti-Spike IgG and IgA, lung resident B cells, and effector memory and resident T cells were efficiently induced, correlating with complete pulmonary protection against the SARS-CoV-2 Delta variant, demonstrating the suitability of the LV::S_Beta-2P_ vaccine candidate as an intranasal booster against COVID-19.

## Introduction

Considering: (i) the sustained pandemicity of coronavirus disease 2019 (COVID-19), (ii) weakening protection potential of the first-generation vaccines against Severe Acute Respiratory Syndrome beta-coronavirus 2 (SARS-CoV-2), and (iii) ceaseless emergence of new viral Variants of Concerns (VOCs), new effective vaccine platforms can be critical for the future primary or booster vaccines [1]. We recently demonstrated the strong performance of a non-integrative lentiviral vaccination vector (LV) encoding the full-length sequence of Spike glycoprotein (S) from the ancestral SARS-CoV-2 (LV::S), when used in systemic prime followed by intranasal (i.n.) boost in multiple preclinical models [2]. LV::S ensures complete (cross) protection of the respiratory tract against ancestral SARS-CoV-2 and VOCs [3]. In addition, in our new transgenic mice expressing human Angiotensin Converting Enzyme 2 (hACE2) and displaying unprecedented permissiveness of the brain to SARS-CoV-2 replication, an i.n. boost with LV::S is required for full protection of the central nervous system [3]. LV::S is intended to be used as a booster for individuals who previously been vaccinated against and/or infected by SARS-CoV-2 to reinforce and broaden protection against emerging VOCs with immune evasion potential [4].

Vaccine LVs are non-integrating, non-replicative, non-cytopathic and negligibly inflammatory [5,6]. These vectors are pseudotyped with the heterologous glycoprotein from Vesicular Stomatitis Virus (VSV-G) which confers them a broad tropism for diverse cell types, including dendritic cells. The latter are mainly non-dividing cells and thus barely permissive to gene transfer. Hence, LVs possess the central property to efficiently transfer genes to the nuclei of not only dividing but also non-dividing cells, which therefore renders possible efficient transduction of non-dividing immature dendritic cells. The resulting endogenous antigen expression in these cells, with unique ability of dendritic cells to activate naïve T cells [7], correlate with a strong ability of LVs at inducing high-quality effector and memory T cells [8]. Importantly, VSV-G pseudo-typing also avoids LVs to be targets of preexisting vector-specific immunity in humans which is key in vaccine development [5,6]. The safety of LV has been established in humans in a phase I/IIa Human Immunodeficiency Virus-1 therapeutic vaccine trial, even though if an integrative version of LV had been used in that clinical trial [9]. Because of their non-cytopathic and non-inflammatory properties [10,11], LVs are well suitable for mucosal vaccination. The i.n. immunization approach is expected to trigger mucosal IgA responses, as well as resident B and T lymphocytes in the respiratory tract [12]. This immunization route has also been shown to be the most effective at reducing SARS-CoV-2 transmission in both hamster and macaque preclinical models [13]. Induction of mucosal immunity by i.n. immunization allows SARS-CoV-2 neutralization, directly at the gateway to the host organism, before it gains access to major infectable anatomical sites [2].

The duration of the protection conferred by the first generation COVID-19 vaccines is not yet well established, hardly predictable with serological laboratory tests and variable in diverse individuals and against distinct VOCs. Despite high vaccination rates, the current exacerbation of the world-wide pandemic indicates that repeated booster immunizations will be needed to ensure individual and collective immunity against COVID-19. In this context, the safety and potential adverse effects of multiple additional homologous doses of the first generation COVID-19 vaccines, for instance related to allergic reaction to polyethylene glycol (PEG) contained in mRNA vaccines, have to be taken into the account [14]. Importantly, an unmatched vaccine delivery method, i.e., heterologous prime-boost format, has been proven to be a more successful strategy than homologous prime-boost approach in numerous preclinical models of various infectious diseases [15–17]. Therefore, new efficient vaccination platforms are of particular interest to develop heterologous boosters against COVID-19. The LV::S vaccine candidate has potential for prophylactic use against COVID-19, mainly based on its strong capacity to induce not only strong neutralizing humoral responses, but also and most importantly, robust protective T-cell responses which preserve their immune detection of spike from SARS-CoV-2 VOCs, despite the accumulation of escape mutations [3]. LV::S is remarkably suitable to be used, as a heterologous i.n. booster vaccine, to reinforce and broaden protection against the emerging VOCs, while collective immunity in early vaccinated nations is waning a few months after completion of the initial immunization, and while new waves of infections are on the rise [4].

In the present study, toward the preparation of a clinical trial, we first generated an LV encoding the down-selected S_CoV-2_ of the Beta variant, stabilized by K^986^P and V^987^P substitutions in the S2 domain of S_CoV-2_ (LV::S_Beta-2P_). In mice, primed and boosted intramuscularly (i.m.) with mRNA vaccine encoding for the ancestral S_CoV-2_ [18,19], and in which the (cross) sero-neutralization potential was progressively decreasing, we investigated the systemic and mucosal immune responses and the protective potential of an i.n. LV::S_Beta-2P_ heterologous boost.

## Results

### Antigen design and down-selection of a lead candidate

To select the most suitable S_CoV-2_ variant to induce the greatest neutralization breadth based on the known variants, we generated LVs encoding the full length S_CoV-2_ from the Alpha, Beta or Gamma SARS-CoV-2 VOCs. C57BL/6 mice (*n* = 5/group) were primed i.m. (wk 0) and boosted i.m. (wk 3) with 1 × 10^8^ TU/mouse of each individual LV and the (cross) neutralization potential of their sera was assessed before boost (wk 3) and after boost (wk 5) against pseudoviruses carrying various S_CoV-2_ **(Figure 1A)**. Immunization with LV::S_Alpha_ generated appropriate neutralization capacity against S_D614G_ and S_Alpha_ but not against S_Beta_ and S_Gamma_ **(Figure 1B)**. Between LV::S_Beta_ and LV::S_Gamma_, the former generated the highest cross sero-neutralization potential against S_D614G_, S_Alpha_ and S_Gamma_ variants. In accordance with previous observations using other vaccination strategies, in the context of immunization with LV, the K^986^P - V^987^P substitutions in the S2 domain of S_CoV-2_ improved the (cross) sero-neutralization potential **(Figure 1C)**, probably due to an extended half-life of S_CoV-2-2P_ [20].

**Figure 1.**
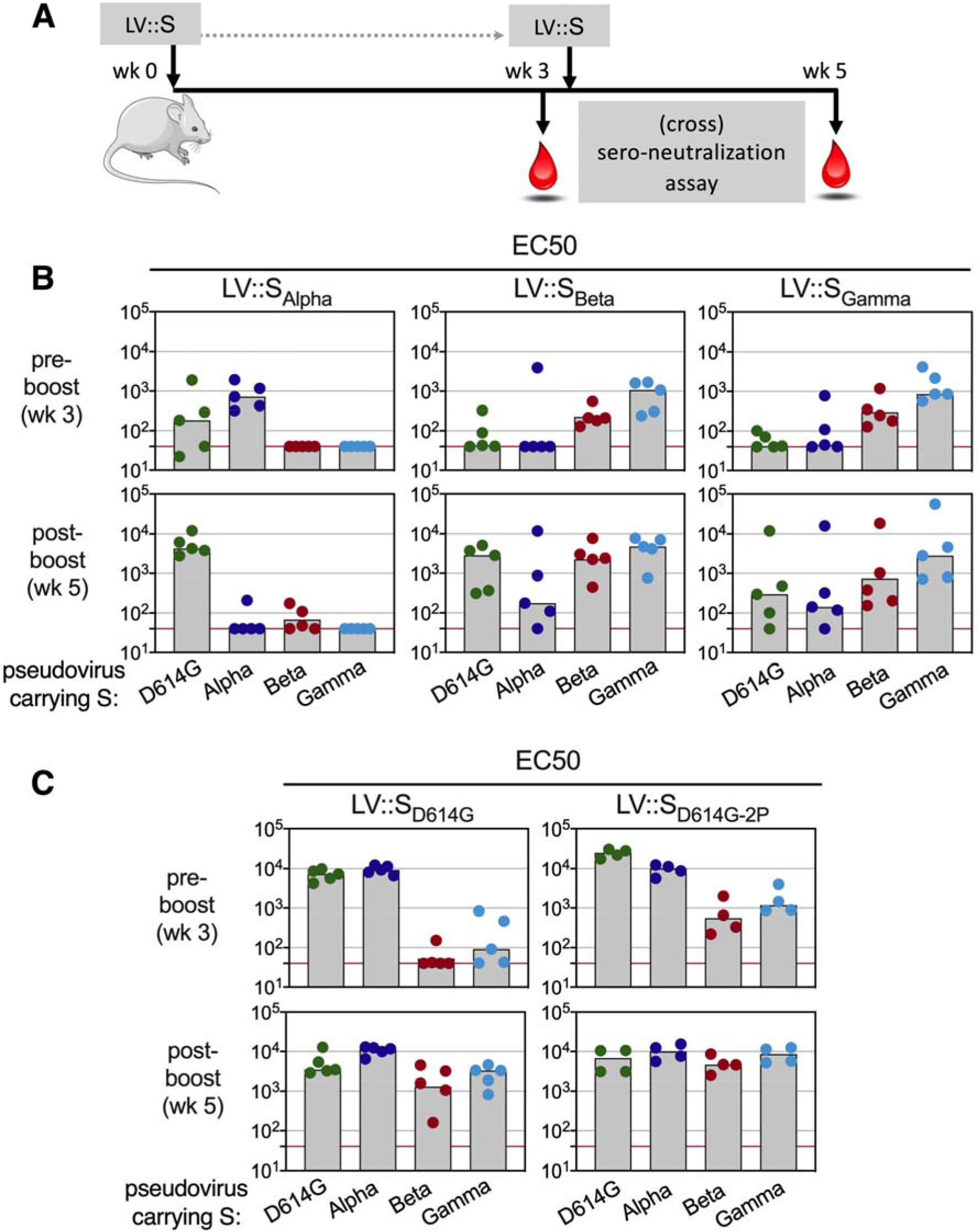
Down-selection of a S_COV-2_ variant with the highest potential to induce cross sero-neutralizing antibodies. **(A)**Timeline of prime-boost vaccination with LV::S_Alpha_, LV::S_Beta_ or LV::S_Gamma_ and (cross) sero-neutralization assays in C57BL/6 mice (*n*= 4-5/group). **(B)** EC50 of neutralizing activity of sera from vaccinated mice was evaluated before and after the boost, against pseudo-viruses carrying S_CoV-2_ from D614G, Alpha, Beta or Gamma variants. **(C)** EC50 of sera from C57BL/6 mice, vaccinated following the regimen detailed in **(A)** with LV encoding for S_D614G_, either WT or carrying the K^986^P - V^987^P substitutions in the S2 domain. EC50 was evaluated before and after the boost, as indicated in **(B)**.

Taken together these data allowed to down select S_Beta-2P_ as the best cross-reactive antigen candidate to be used in the context of LV (LV::S_Beta-2P_) to strengthen the waning immunity previously induced by the first generation COVID-19 vaccines, like mRNA.

### Follow-up of humoral immunity in mRNA-primed and -boosted mice and effect of LV::S_Beta-2P_ i.n. boost

We analyzed the potential of LV::S_Beta-2P_ i.n. boost vaccination to strengthen and broaden the immune responses in mice which were initially primed and boosted with mRNA and in which the (cross) sero-neutralization potential was decreasing. C57BL/6 mice were primed i.m. at wk 0 and boosted i.m. at wk 3 with 1 μg/mouse of mRNA **(Figure 2A)**. In mRNA-primed mice, serum anti-S_CoV-2_ and anti-RBD IgG were detected at wk 3, increased after mRNA boost as studied at wk 6 and 10, and then decreased at wk 17 in the absence of an additional boost **(Figure S1A)**.

**Figure 2.**
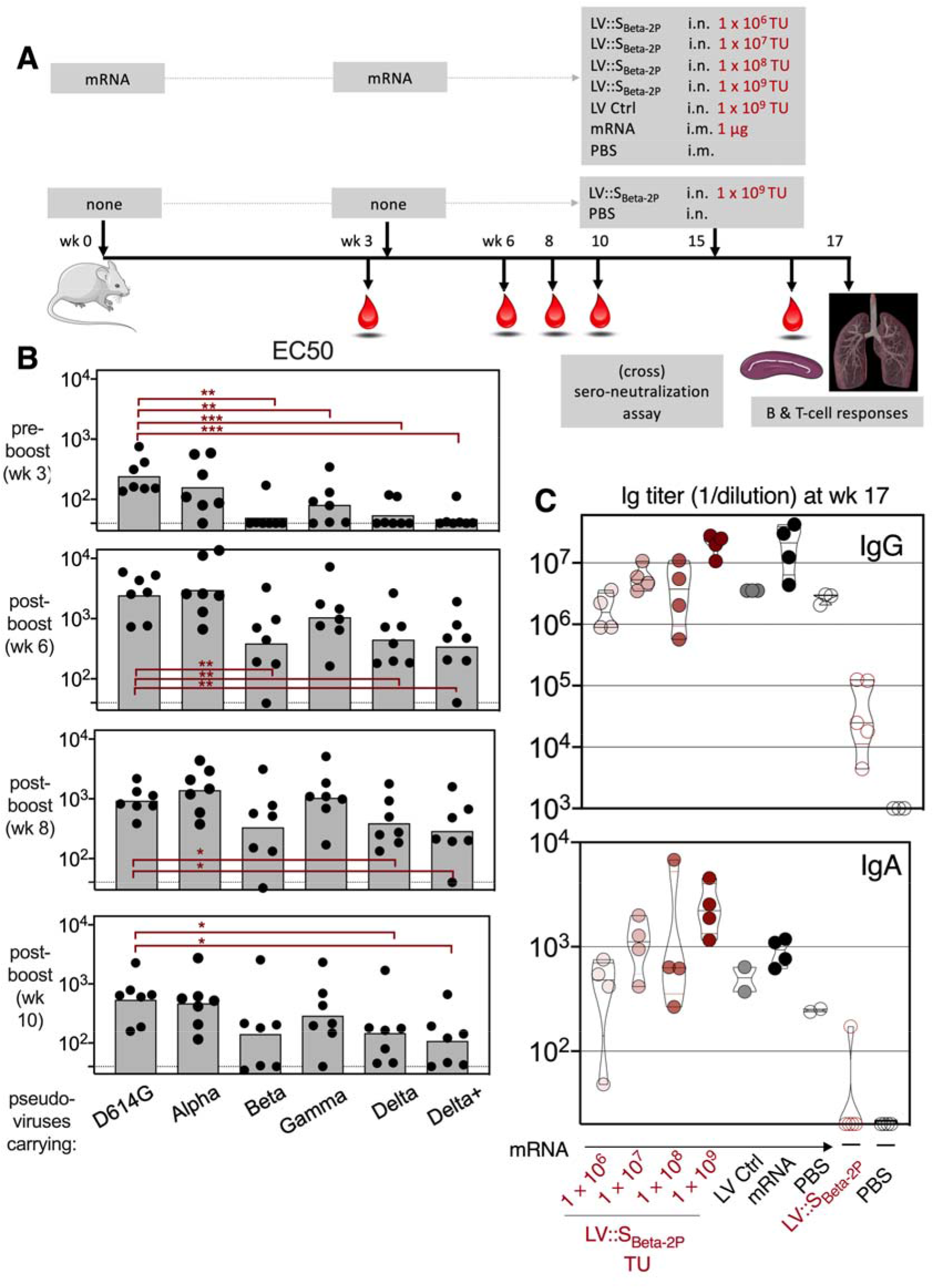
Anti-S_CoV-2_ humoral responses in mRNA-vaccinated mice which were further intranasally boosted with LV::S_Beta-2P_. **(A)**Timeline of mRNA i.m.-i.m. prime-boost vaccination in C57BL/6 mice which were later immunized i.n. by escalating doses of LV::S_Beta-2P_ (*n* = 4-5/group) and the (cross) sero-neutralization follow-up. **(B)**Serum EC50 determined at the indicated time points against pseudo-viruses carrying S_CoV-2_ from D614G, Alpha, Beta, Gamma, Delta or Delta+ variants. **(C)** Anti-S_CoV-2_ IgG (upper panel) or IgA (lower panel) titers in the sera two weeks after i.n. LV::S_Beta-2P_ boost. Statistical significance was determined by Mann-Whitney test (*=*p*< 0.05, **=*p*< 0.01, ***=*p*< 0.001).

Longitudinal serological follow-up demonstrated that at 3 wks post prime, cross-neutralization activities against both S_D614G_ and S_Alpha_ were readily detectable **(Figure 2B)**. Cross sero-neutralization was also detectable, although to a lesser degree, against S_Gamma_, but not against S_Beta_, S_Delta_ or S_Delta+_. At wk 6, i.e., 3 wks post boost, cross sero-neutralization activities against all S_CoV-2_ variants were detectable, although at significantly lesser extents against S_Beta_, S_Delta_ and S_Delta+_. From wk 6 to wk 10, cross sero-neutralization against S_Beta_, S_Delta_, or S_Delta+_gradually and significantly decreased. At wk 10, half of the mice lost the cross sero-neutralization potential against S_Beta_, S_Delta_, or S_Delta+_ **(Figure 2B)**.

At wk 15, groups of mRNA-primed and -boosted mice received i.n. escalating doses of 1 ×10^6^, 1 × 10^7^, 1 × 10^8^, or 1 × 10^9^ Transduction Units (TU)/mouse of LV::S_Beta-2P_ **(Figure 2A)**. Control mRNA-primed and -boosted mice received i.n. 1 × 10^9^ TU of an empty LV (LV Ctrl). In parallel, at this time point, mRNA-primed and -boosted mice were injected i.m. with 1 μg of mRNA-vaccine or PBS. The dose of 1 μg of mRNA per mouse has been demonstrated to be fully protective in mice [21]. Unprimed, age-matched mice received i.n. 1 × 10^9^ TU of LV::S_Beta-2P_ or PBS.

In the previously mRNA-primed and -boosted mice, injected at wk 15 with 1 × 10^8^ or 1 × 10^9^ TU of LV::S_Beta-2P_ or a third dose of mRNA, marked anti-S_CoV-2_ IgG titer increases were observed **(Figure 2C)**. The titers of anti-S_CoV-2_ IgA were higher in the mice injected with 1 × 10^9^ TU of LV::S_Beta-2P_ than those injected with a third 1 μg dose of mRNA vaccine **(Figure 2C)**. At the mucosal level, at this time point, titers of anti-S_CoV-2_ and anti-RBD IgG in the total lung extracts increased in a dose-dependent manner in LV::S_Beta-2P_-boosted mice, and the titer obtained with the highest dose of LV::S_Beta-2P_ was comparable to that after the third 1 μg i.m. dose of mRNA vaccine **(Figure S1B)**. Importantly, significant titers of lung anti-S_CoV-2_ IgA were only detected in LV::S_Beta-2P_-boosted mice **(Figure S1B)**.

At the lung cellular level, CD19^+^ B cells which are class-switched and thus surface IgM^-^/IgD^-^plasma cells, and which express CD38, CD62L, CD73 and CD80, can be defined as lung resident B cells (Brm) [22,23] **(Figure 3A)**. The proportion of these B cells increased in a dose-dependent manner in the lungs of mice boosted i.n. with LV::S_Beta-2P_ **(Figure 3B)**. Mucosal anti-S_CoV-2_ IgA and Brm were barely detectable in the mice boosted i.m. at wk15 with 1 μg of mRNA, which was the single dose of RNA tested in our experiment and thus serves only as indication.

**Figure 3.**
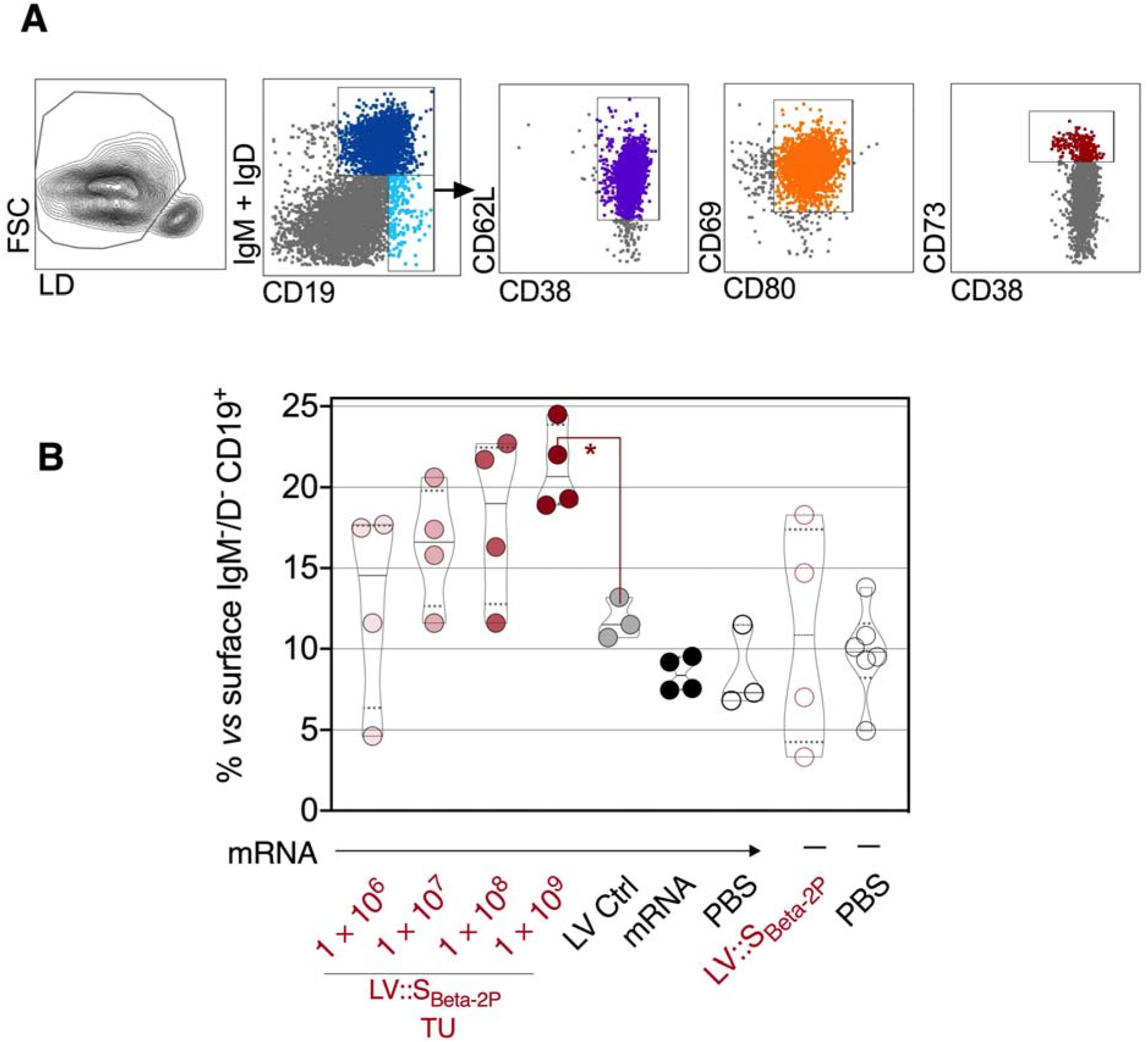
Lung B-cell resident memory subset in mRNA-vaccinated mice which were further intranasally boosted with LV::S_Beta-2P_. The mice are those detailed in the Figure 2. Mucosal immune cells were studied two weeks after LV::S_Beta-2P_ i.n. boost. **(A)** Cytometric gating strategy to detect lung Brm in mRNA-vaccinated mice which were further intranasally boosted with LV::S_Beta-2P_. **(B)**Percentages of these cells among lung CD19^+^ surface IgM^-^/IgD^-^ B cells in mRNA-vaccinated mice which were further intranasally boosted with LV:: S_Beta-2P_. Statistical significance was determined by Mann-Whitney test (*=*p*< 0.05).

### Systemic and mucosal T-cell immunity after i.n. LV::S_Beta-2P_ boost in previously mRNA-primed and -boosted mice

Mice were primed and boosted with 1 μg mRNA-vaccine and then boosted i.n. at wk15 with escalating doses of LV::S_Beta-2P_, according to the above-mentioned regimen **(Figure 2A)**. At wk 17, i.e., two wks after the late boost, systemic anti-S_CoV-2_ T-cell immunity was assessed by IFN-γ-specific ELISPOT in the spleen of individual mice after in vitro stimulation with individual S:256-275, S:536-550 or S:576-590 peptide, encompassing immunodominant S_CoV-2_ regions for CD8^+^ T cells in H-2^b^ mice [2]. Importantly, the weak anti-S CD8^+^ T-cell immunity, detectable in the spleen of mRNA-primed-boosted mice at wk 17, largely increased following i.n. boost with 1 × 10^8^ and 1 × 10^9^ TU of LV::S_Beta-2P_, similarly to the increase after i.m. mRNA boost **(Figure 4)**.

**Figure 4.**
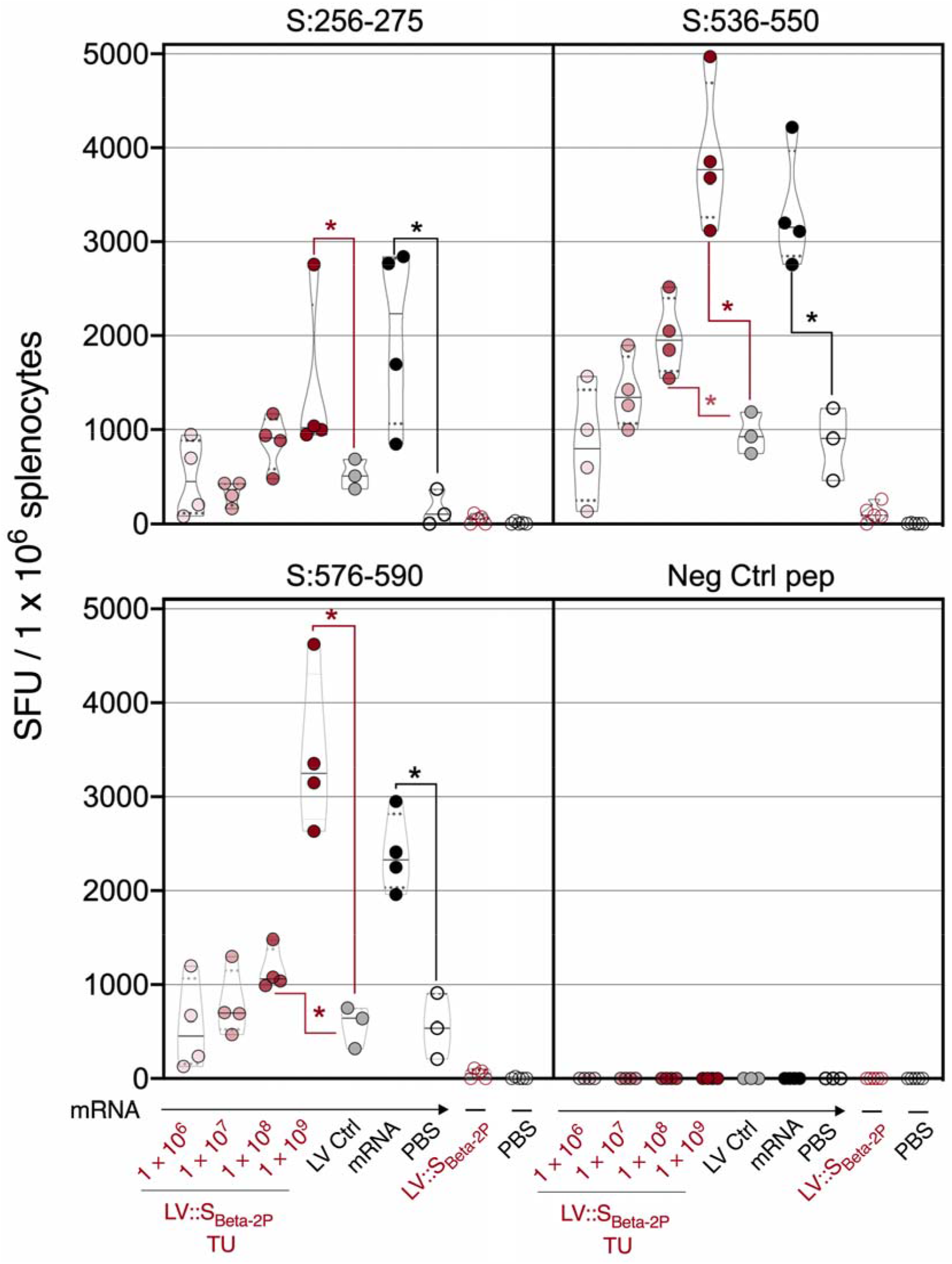
Systemic CD8^+^ T-cell responses to S_COV-2_ in mRNA-vaccinated mice which were further intranasally boosted with LV::S_Beta-2P_. The mice are those detailed in the Figure 2. T-splenocyte responses were evaluated two weeks after LV::S_Beta-2P_ i.n. boost by IFN-γ ELISPOT after stimulation with S:256-275, S:536-550 or S:576-590 synthetic 15-mer peptides encompassing S_CoV-2_ MHC-I-restricted epitopes. Statistical significance was evaluated by Mann-Whitney test (*=*p*< 0.05).

In parallel, in the same animals, the mucosal anti-S_CoV-2_ T-cell immunity was assessed by intracellular Tc1 and Tc2 cytokine staining in T cell-enriched fraction from individual mice after in vitro stimulation with autologous bone-marrow-dendritic cells loaded with a pool of S:256-275, S:536-550 and S:576-590 peptides **(Figure 5)**. In previously mRNA-primed and -boosted mice, only a few S_CoV-2_-specific IFN-γ/TNF/IL-2 CD8^+^ T-cell responses were detected in the lungs **(Figure 5)**. The i.n. administration of LV::S_Beta-2P_ boosted, in a dose dependent manner, these Tc1 responses. Sizable percentages of these Tc1 cells were induced with 1 × 10^8^ or 1 × 10^9^ TU of LV::S_Beta-2P_. mRNA (1 μg) i.m. administration had a substantially lower boost effect on mucosal T cells **(Figure 5)**. Tc2 responses (IL-4, IL-5, IL-10 and IL-13) were not detected in any experimental group **(Figure S2)**, as assessed in the same lung T-cell cultures.

**Figure 5.**
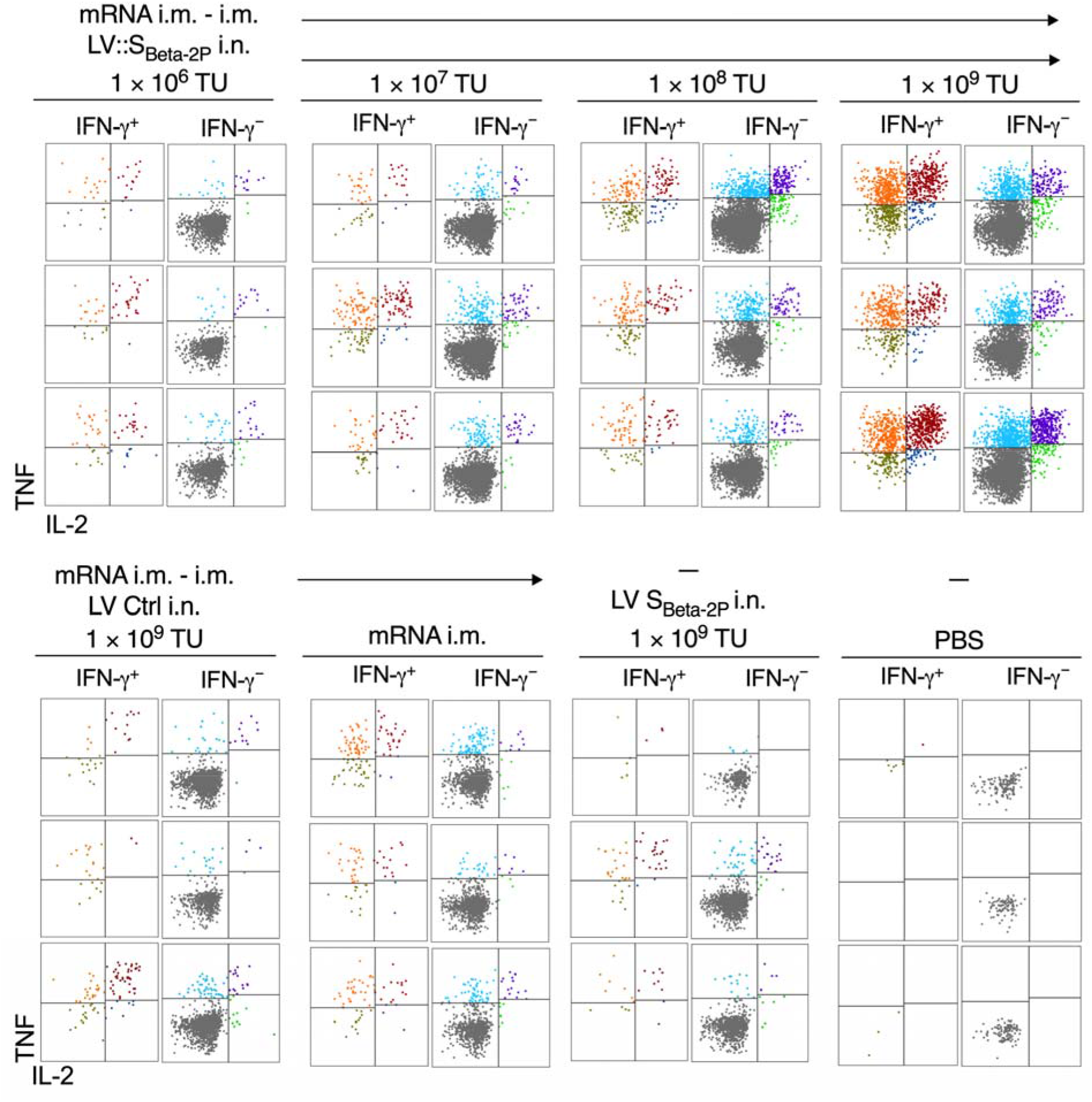
Mucosal CD8^+^ T-cell responses to S_COV-2_ in mRNA-vaccinated mice which were further intranasally boosted with LV::S_Beta-2P_. The mice are those detailed in the Figure 2. **(A)** Representative IFN-γ response by lung CD8^+^ T cells detected by intracellular cytokine staining after in vitro stimulation with a pool of S:256-275, S:536-550 and S:576-590 peptides. Cells are gated on alive CD45^+^ CD8^+^ T cells.

Mucosal lung resident memory T cells (Trm), CD8^+^ CD44^+^ CD69^+^ CD103^+^ which are one of the best correlates of protection in the infectious diseases [24], were readily detected in the mice boosted i.n. with 1 × 10^8^ or 1 × 10^9^ TU of LV::S_Beta-2P_ **(Figure 6A, B)**. No Trm were detected in the lungs of mice boosted late with 1 μg mRNA i.m..

**Figure 6.**
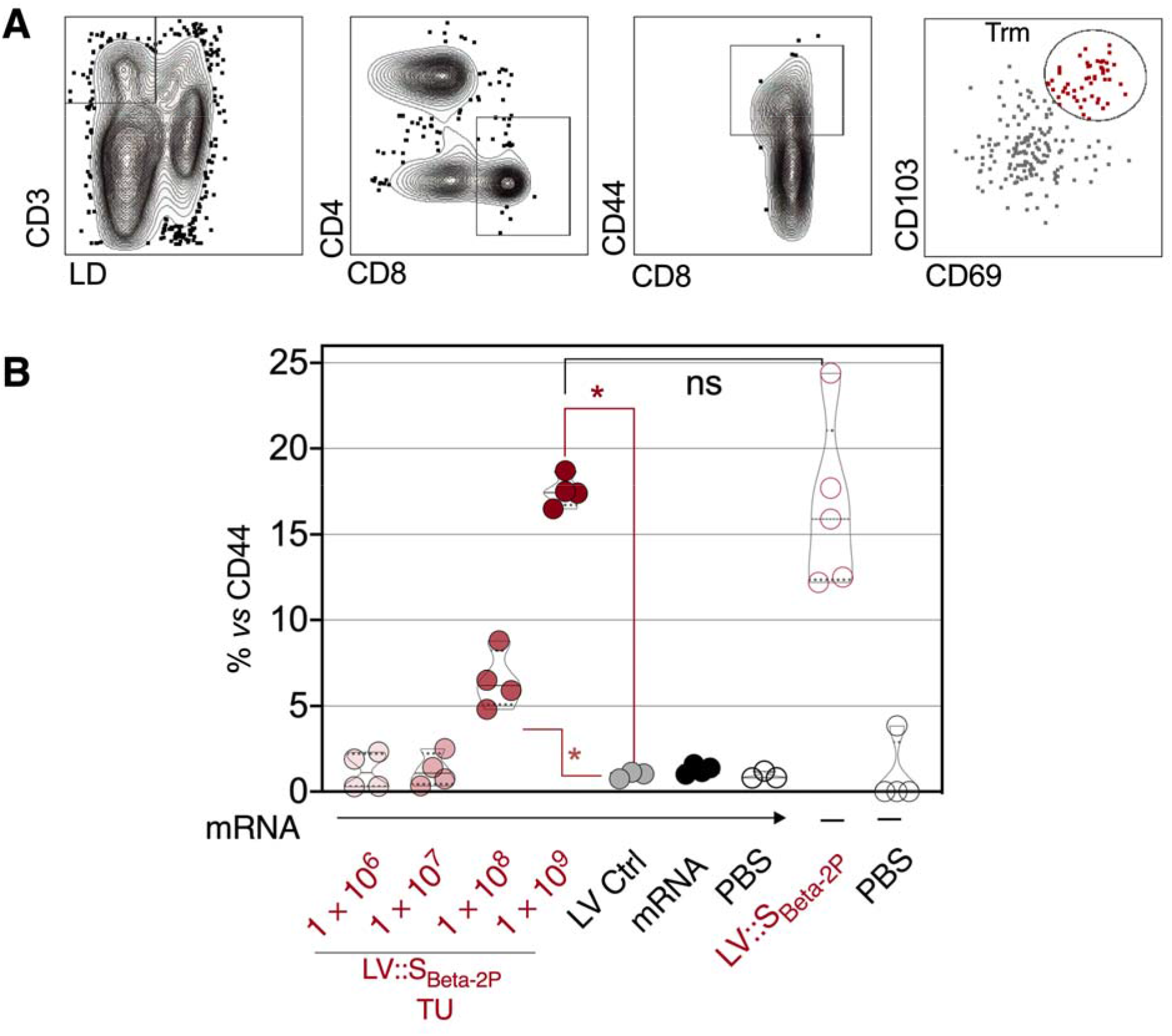
Lung T resident memory subset in mRNA-vaccinated mice which were further intranasally boosted with LV::S_Beta-2P_. The mice are those detailed in the Figure 2. Mucosal immune cells were studied two weeks after LV::S_Beta-2P_ i.n. boost. **(A)** Cytometric gating strategy to detect lung CD8^+^ T resident memory (CD44^+^CD69^+^CD103^+^), and **(B)** percentages of this subset among CD8^+^ CD44^+^ T-cells in mRNA-vaccinated mice which were further intranasally boosted with LV::S_Beta-2P_. Statistical significance was evaluated by Mann-Whitney test (*=*p*< 0.05).

### Features of lungs after LV::S_Beta-2P_ i.n. administration

To identify the immune cell subsets transduced in vivo by LV after i.n. administration, C57BL/6 mice were immunized i.n. with the high dose of 1 □ × □ 10^9^ TU of LV::GFP or LV::nano-Luciferase (LV::nLuc) as a negative control. Lungs were collected at 4 □ days post-immunization and analyzed by cytometry in individual mice. CD45^-^ cell subset was devoid of GFP^+^ cells. Only a very few GFP^+^ cells were detected in the CD45^+^ hematopoietic cells **(Figure S3A-C)**. The CD45^+^ GFP^+^ cells were located in a CD11b^hi^ subset and in the CD11b^int^ CD11c^+^ CD103^+^ MHC-II^+^(dendritic cells) **(Figure S3B)**.

To evaluate possible lung infiltration after LV i.n. administration, C57BL/6 mice were injected i.n. with the high dose of 1 □ × □ 10^9^ TU of LV::S_Beta-2P_ or PBS as a negative control. Lungs were collected at 1, 3□or 14 days post-injection for histopathological analysis. H&E histological sections displayed minimal to moderate inflammation, interstitial and alveolar syndromes in both experimental groups, regardless of the three time points investigated. No specific immune infiltration or syndrome was detected in the animals treated i.n. with 1 □ × □ 10^9^ TU of LV::S_Beta-2P_, compared to PBS **(Figure S4A, B)**.

### Protection of lungs in mRNA-primed and -boosted mice, and later boosted i.n. with LV::S_Beta-2P_

We then evaluated the protective vaccine efficacy of LV::S_Beta-2P_ i.n. in mRNA-primed and - boosted mice. At wk 15, mRNA-primed and -boosted mice received i.n. 1 × 10^8^ TU of LV::S_Beta-2P_or control empty LV. **(Figure 7A)**. The choice of this dose was based on our previous experience with this dose which was effective in protection in homologous LV::S prime-boost regimens [2,3]. Control mRNA-immunized mice received i.m. 1 μg of mRNA or PBS. Unvaccinated, age-and sex-matched controls were left unimmunized. Five weeks after the late boost, i.e. at wk 20, all mice were pre-treated with 3 × 10^8^ Infectious Genome Units (IGU) of an adenoviral vector serotype 5 encoding hACE2 (Ad5::hACE2) [2] to render their lungs permissive to SARS-CoV-2 replication **(Figure 7A)**. Four days later, mice were challenged with SARS-CoV-2 Delta variant, which at the time of this study, i.e. November 2021, was the dominant SARS-CoV-2 variant worldwide.

**Figure 7.**
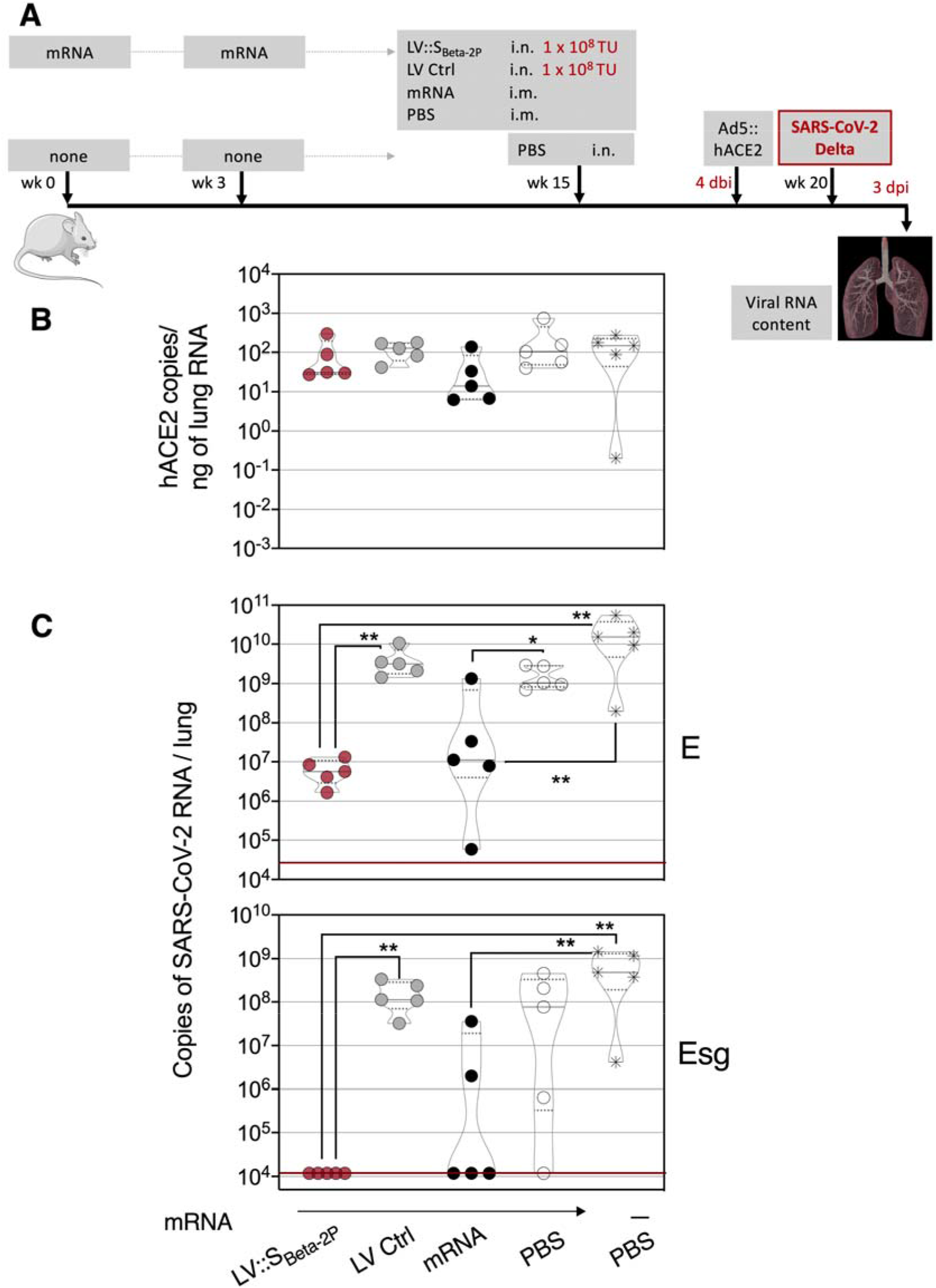
Full protective capacity of LV::S_Beta-2P_ i.n. boost against Delta variant in initially mRNA-primed and boosted mice. **(A)** Timeline of mRNA i.m.-i.m. prime-boost vaccination in C57BL/6 mice which were later immunized i.n. with 1 × 10_8_ TU/mouse of LV::S_Beta-2P_(*n*= 4-5/group), pre-treated i.n. with Ad5::hACE-2 4 days before i.n. challenge with 0.3 × 10^5^ TCID50 of SARS-CoV-2 Delta variant. **(B)** Comparative quantification of *hACE-2* mRNA in the lungs of Ad5::hACE-2 pre-treated mice at 3 dpi. **(C)** Lung viral RNA contents, evaluated by conventional Especific (top) or sub-genomic Esg-specific (bottom) qRT-PCR at 3 dpi. Red lines indicate the detection limits. Statistical significance was evaluated by Mann-Whitney test (*=*p* < 0.05, **=*p* < 0.01).

At day 3 post infection, primary analysis of the total lung RNA showed that hACE2 mRNA was similarly expressed in all mice after Ad5::hACE2 *in vivo* transduction **(Figure 7B)**. Lung viral loads were then determined at 3 dpi by assessing total E RNA and sub-genomic (Esg) E_CoV-2_ RNA qRT-PCR, the latter being an indicator of active viral replication [25–27]. In mice initially primed and boosted with mRNA-vaccine and then injected i.n. with the control LV or i.m. with PBS, no significant protective capacity was detectable **(Figure 7C)**. In contrast, the LV::S_Beta-2P_ i.n. boost of the initially mRNA vaccinated mice drastically reduced the total E RNA content of SARS-CoV-2 and no copies of the replication-related Esg E_CoV-2_ RNA were detected in this group **(Figure 7C)**. In the group which received a late mRNA i.m. boost, the total E RNA content was also significantly reduced and the content of Esg E_CoV-2_ RNA was undetectable in 3 out of 5 in this group.

Therefore, a late LV::S_Beta-2P_ heterologous i.n. boost, given at wk 15 after the first injection of mRNA, at the dose of 1 × 10^8^ TU/mouse resulted in complete protection, i.e. total absence of viral replication in 100% of animals, against a high dose challenge with SARS-CoV-2 Delta variant.

## Discussion

With the weakly persistent prophylactic potential of the immunity initially induced by the first-generation COVID-19 vaccines, especially against new VOCs, administration of additional vaccine doses becomes essential [1]. As an alternative to additional doses of the same vaccines, combining vaccine platforms in a heterologous prime-boost regimen holds promise for gaining protective efficacy [28]. Compared to homologous vaccine dose administrations, heterologous prime-boost strategies may reinforce more efficiently specific adaptive immune responses and long-term protection [29]. Furthermore, the sequence of the Spike antigen has to be adapted according to the dynamics of SARS-CoV-2 VOC emergence in order to induce the greatest neutralization breadth. Protection against symptomatic SARS-CoV-2 infection is mainly related to sero-neutralizing activity, while CD8^+^ T-cell immunity, with their ability to cytolyze virus-infected cells, especially control the virus replication and result in resolution of viral infection [30]. Therefore, an appropriate B- and T-cell vaccine platform, including an adapted S_CoV2_ sequence, is of utmost interest at the current step of the pandemic.

The LV-based strategy, which is highly efficient, not only in inducing humoral responses but also, and particularly, in establishing high quality and memory T-cell responses [8], is a favorable platform for a heterologous boost, even if it is also largely efficacious by its own as a primary COVID-19 vaccine candidate [2,3]. Furthermore, LV is non-cytopathic, non-replicative and scarcely inflammatory and thus can be used to perform non-invasive i.n. boost to efficiently induce sterilizing mucosal immunity, which protects the respiratory system as well as the central nervous system [2,3]. The i.n. route of vaccination has been shown by several teams to be the most efficacious route at reducing viral contents in nasal swabs and nasal olfactory neuroepithelium [31,32], which can contribute to block the respiratory chain of SARS-CoV-2 transmission. One of the advantages of LV-based immunization is the induction of strong T-cell immune responses with high cross-reactivity of T-cell epitopes from Spike of diverse VOCs. Therefore, when the neutralizing antibody fails or wanes, the T-cell arm remains largely protective, as we recently described in antibody-deficient, B-cell compromised μMT KO mice [3]. This property is relative to a high-quality and long-lasting T-cell immunity induced against multiple preserved T-cell epitopes, despite the mutations accumulated in the Spike of the emerging VOCs [3], including the Omicron variant.

In the present study, we down-selected the S_Beta-2P_ antigen which induced the greatest neutralization breadth against the main SARS-CoV-2 VOCs and designed a non-integrative LV encoding a stabilized version of this antigen. In mice primed and boosted with mRNA vaccine (encoding the ancestral S_CoV-2_ sequence), with waning (cross) sero-neutralization capacity, we used escalating doses of LV::S_Beta-2P_ for an i.n. late boost. We demonstrated a dose-dependent increase in anti-S_CoV-2_ IgG and IgA titers, and a broadened sero-neutralization potential both in the sera and lung homogenates against VOCs. No anti-S_CoV-2_ IgA was detected in the lungs of mice injected with the third dose of 1 μg of mRNA given via i.m.. Increasing proportions of non-circulating Brm, defined as class-switched surface IgM^-^/IgD^-^ plasma cells, with CD38^+^ CD73^+^ CD62L^+^ CD69^+^CD80^+^ phenotype [22,23], were detected in a dose-dependent manner, in the lungs of mice boosted i.n. with LV::S_Beta-2P_.

Spike-specific, effector lung CD8^+^ Tc1 cells were largely detected in the initially mRNA-primed and boosted mice which received a late i.n. LV::S_Beta-2P_ boost. These lung CD8^+^ T cells did not display Tc2 phenotype. Increasing proportions of lung CD8^+^ CD44^+^ CD69^+^ CD103^+^ Trm were also detected, in a dose-dependent manner, only in LV::S_Beta-2p_ i.n. boosted mice. The systemic CD8^+^ T-cell responses against various immunogenic regions of S_CoV-2_ were also increased with 1 × 10^8^ or 1 × 10^9^ TU doses of LV::S_Beta-2P_ i.n. boost in the initially mRNA-primed and -boosted mice. The highest i.n. dose of of LV::S_Beta-2P_ was comparable to the third injection of 1 μg/mouse of mRNA given by i.m.. The fact that the i.n. administration of LV::S_Beta-2P_ had a substantial boost effect on the systemic T-cell immunity indicates that this boost pathway is not at the expense of the induction of systemic immunity. Evaluation of the protection in mRNA-primed and -boosted mice, showed that 20 wk after the first injection of mRNA vaccine, there was no protection detectable in the lungs against infection with the SARS-CoV-2 Delta variant. Importantly, an i.n. boost at wk 15 with the dose of 1 × 10^8^ TU/mouse LV::S_Beta-2P_ resulted in full inhibition of SARS-CoV-2 replication in the lungs upon challenge with the Delta variant at wk 20. In the mice receiving an 1 μg mRNA i.m. boost at wk 15 the lung SARS-CoV-2 RNA contents was reduced in a statistically comparable manner, albeit without total inhibition of viral replication in all mice.

The lack of protection against the Delta variant infection only four months after the initial systemic prime-boost by mRNA vaccine, as we observed in the preclinical model in this study, may be explained by the weak efficiency of the ancestral S_CoV-2_ sequence to induce long-lasting neutralizing antibodies against the recent VOCs. In addition, it can be hypothesized that the adaptive immune memory induced by i.m. mRNA immunization is likely to be localized in secondary lymphoid organs at anatomical sites located far from the upper respiratory tract. In such a context, the extraordinary rapid replication of new VOCs, such as Delta or Omicron, in the upper respiratory tract would not leave enough time for the reactivation of immune memory from remote anatomical sites and the recruitment of the immune arsenal from these sites. In human populations, such scenario would lead to a high possibility of viral replication and variable levels of its transmission, which would prevent the epidemic from being completely contained by mass vaccination through the systemic route.

It is not yet known if a single third booster will extend and maintain the protective potential, or whether semi-regular boosters will be required against COVID-19 in the future. The LV::S_Beta-2P_ i.n. boost strengthens the intensity, broadens the VOC cross-recognition, and targets B-and T-cell immune responses to the principal entry point of SARS-CoV-2 to the mucosal respiratory tract of the host organism and avoid the infection of main anatomical sites. It is interesting to note that in rodents only a pair of cervical ganglia exists versus a large network of such ganglia in humans [33]. Following i.n. immunization, this anatomical feature in humans may provide even a more consistent site of immune response induction and local memory maintenance, at the vicinity of to the potential site of airway infection. In addition, nasopharynx-associated lymphoid tissue is a powerful defense system composed of: (i) organized lymphoid tissue, i.e., tonsils, and (ii) a diffuse nose-associated lymphoid tissues, where effector and memory B and T lymphocytes are able to maintain long-lasting immunity [34]. This mucosal immune arsenal deserves to be explored in the control of SARS-CoV-2 transmission in the current context of the pandemic. A phase I clinical trial is currently in preparation for the use of i.n. boost by LV::S_Beta-2P_ in previously vaccinated humans or in COVID-19 convalescents.

## Materials and Methods

### Mice immunization and SARS-CoV-2 infection

Female C57BL/6JRj mice were purchased from Janvier (Le Genest Saint Isle, France), housed in individually-ventilated cages under specific pathogen-free conditions at the Institut Pasteur animal facilities and used at the age of 7 wks. Mice were immunized i.m. with 1 μg/mouse of mRNA-1273 (Moderna) vaccine. For i.n. injections with LV, mice were anesthetized by i.p. injection of Ketamine (Imalgene, 80 mg/kg) and Xylazine (Rompun, 5 mg/kg). For protection experiments against SARS-CoV-2, mice were transferred into filtered cages in isolator. Four days before SARS-CoV-2 inoculation, mice were pretreated with 3 × 10^8^ IGU of Ad5::hACE2 as previously described [2]. Mice were then transferred into a level 3 biosafety cabinet and inoculated i.n. with 0.3 × 10^5^ TCID_50_ of the Delta variant of SARS-CoV-2 clinical isolate [35] contained in 20 μl. Mice were then housed in filtered cages in an isolator in BioSafety Level 3 animal facilities. The organs recovered from the infected animals were manipulated according to the approved standard procedures of these facilities.

### Ethical approval of animal experimentation

Experimentation on animals was performed in accordance with the European and French guidelines (Directive 86/609/CEE and Decree 87-848 of 19 October 1987) subsequent to approval by the Institut Pasteur Safety, Animal Care and Use Committee, protocol agreement delivered by local ethical committee (CETEA #DAP20007, CETEA #DAP200058) and Ministry of High Education and Research APAFIS#24627-2020031117362508 v1, APAFIS#28755-2020122110238379 v1.

### Construction and production of vaccinal LV::S_Beta-2P_

First, a codon-optimized sequence of Spike from the Ancestral, D614G, Alpha, Beta or Gamma VOCs were synthetized and inserted into the pMK-RQ_S-2019-nCoV_S501YV2 plasmid. The S sequence was then extracted by BamHI/XhoI digestion to be ligated into the pFlap lentiviral plasmid between the BamHI and XhoI restriction sites, located between the native human ieCMV promoter and the mutated *atg* starting codon of Woodchuck Posttranscriptional Regulatory Element (WPRE) sequence (**Figure S5**). To introduce the K^986^P-V^987^P “2P”double mutation in S_D614G_ or S_Beta_, a directed mutagenesis was performed by use of Takara In-Fusion kit on the corresponding pFlap plasmids. Various pFlap-ieCMV-S-WPREm or pFlap-ieCMV-S2P-WPREm plasmids were amplified and used to produce non-integrative vaccinal LV, as described elsewhere [2,6].

### Analysis of humoral and systemic T-cell immunity

Anti-S_CoV-2_ IgG and IgA antibody titers were determined by ELISA by use of recombinant stabilized S_CoV-2_ or RBD fragment for coating. Neutralization potential of clarified and decomplemented sera or lung homogenates was quantitated by use of lentiviral particles pseudo-typed with S_CoV-2_ from diverse variants, as previously described [2,36].

T-splenocyte responses were quantitated by IFN-γ ELISPOT after in vitro stimulation with S:256-275, S:536-550 or S:576-590 synthetic 15-mer peptides which contain S_CoV-2_ MHC-I-restricted epitopes in H-2^d^ mice [2]. Spots were quantified in a CTL Immunospot S6 ultimate-V Analyser by use of CTL Immunocapture 7.0.8.1 program.

### Phenotypic and Functional cytometric analysis of lung immune cells

Enrichment and staining of lung immune cells were performed as detailed elsewhere [2,3] after treatment with 400 U/ml type IV collagenase and DNase I (Roche) for a 30-minute incubation at 37°C and homogenization by use of GentleMacs (Miltenyi Biotech). Cell suspensions were then filtered through 100 μm-pore filters, centrifuged at 1200 rpm and enriched on Ficoll gradient after 20 min centrifugation at 3000 rpm at RT, without brakes. The recovered cells were co-cultured with syngeneic bone-marrow derived dendritic cells loaded with a pool of A, B, C peptides, each at 1 μg/ml or negative control peptide at x μg/ml. The following mixture was used to detect lung Tc1 cells: PerCP-Cy5.5-anti-CD3 (45-0031-82, eBioScience), eF450-anti-CD4 (48-0042-82, eBioScience) and APC-anti-CD8 (17-0081-82, eBioScience) for surface staining and BV650-anti-IFN-g (563854, BD), FITC-anti-TNF (554418, BD) and PE-anti-IL-2 (561061, BD) for intracellular staining. The following mixture was used to detect lung Tc2 cells: PerCP-Cy5.5-anti-CD3 (45-0031-82, eBioScience), eF450-anti-CD4 (48-0042-82, eBioScience), BV711-anti-CD8 (563046, BD Biosciences), for surface staining and BV605-anti-IL-4 (504125, BioLegend Europe BV), APC-anti-IL-5 (504306, BioLegend Europe BV), FITC-anti-IL-10 (505006, BioLegend Europe BV), PE-anti-IL-13 (12-7133-81, eBioScience) for intracellular staining. The intracellular staining was performed by use of the Fix Perm kit (BD), following the manufacturer’s protocol. Dead cells were excluded by use of Near IR Live/Dead (Invitrogen). Staining was performed in the presence of FcγII/III receptor blocking anti-CD16/CD32 (BD).

To identify lung resident memory CD8^+^ T-cell subsets, a mixture of PerCP-Vio700-anti-CD3 (130-119-656, Miltenyi Biotec), PECy7-CD4 (552775, BD Biosciences), BV510-anti-CD8 (100752, BioLegend), PE-anti-CD62L (553151, BD Biosciences), APC-anti-CD69 (560689, BD Biosciences), APC-Cy7-anti-CD44 (560568, BD Biosciences), FITC-anti-CD103 (11-1031-82, eBiosciences) and yellow Live/Dead (Invitrogen) was used. Lung Brm were studied by surface staining with a mixture of PerCP Vio700-anti-IgM (130-106-012, Miltenyi), and PerCP Vio700-anti-IgD (130-103-797, Miltenyi), APC-H7-anti-CD19 (560143, BD Biosciences), PE-anti-CD38 (102708, BioLegend Europe BV), PE-Cy7-anti-CD62L (ab25569, AbCam), BV711-anti-CD69 (740664, BD Biosciences), BV421-anti-CD73 (127217, BioLegend Europe BV), FITC-anti-CD80 (104705, BioLegend Europe BV and yellow Live/Dead (Invitrogen).

Cells were incubated with appropriate mixtures for 25 minutes at 4°C, washed in PBS containing 3% FCS and fixed with Paraformaldehyde 4% after an overnight incubation at 4°C. Samples were acquired in an Attune NxT cytometer (Invitrogen) and data analyzed by FlowJo software (Treestar, OR, USA).

### Determination of viral RNA content in the organs

Organs from mice were removed and immediately frozen at −80°C on dry ice. RNA from circulating SARS-CoV-2 was prepared from lungs as described elsewhere [2]. Lung homogenates were prepared by thawing and homogenizing in lysing matrix M (MP Biomedical) with 500 μl of PBS using a MP Biomedical Fastprep 24 Tissue Homogenizer. RNA was extracted from the supernatants of organe homogenates centrifuged during 10 min at 2000g, using the Qiagen Rneasy kit, except that the neutralization step with AVL buffer/carrier RNA was omitted. The RNA samples were then used to determine viral RNA content by E-specific qRT-PCR. To determine viral RNA content by Esg-specific qRT-PCR, total RNA was prepared using lysing matrix D (MP Biomedical) containing 1 mL of TRIzol reagent (ThermoFisher) and homogenization at 30 s at 6.0 m/s twice using MP Biomedical Fastprep 24 Tissue Homogenizer. The quality of RNA samples was assessed by use of a Bioanalyzer 2100 (Agilent Technologies). Viral RNA contents were quantitated using a NanoDrop Spectrophotometer (Thermo Scientific NanoDrop). The RNA Integrity Number (RIN) was 7.5-10.0. SARS-CoV-2 E or E sub-genomic mRNA were quantitated following reverse transcription and real-time quantitative TaqMan^®^ PCR, using SuperScriptTM III Platinum One-Step qRT-PCR System (Invitrogen) and specific primers and probe (Eurofins), as recently described [3].

### Lung histology

Left lobes from lungs were fixed in formalin and embedded in paraffin. Paraffin sections (5-μm thick) were stained with Hematoxylin and Eosin (H&E). Slides were scanned using the AxioScan Z1 (Zeiss) system and images were analyzed with the Zen 2.6 software. Histological images were evaluated according to a score of 0 to 5 (normal, minimal, mild, moderate, marked, severe).

## Acknowledgments

The authors are grateful to Dr. Marie José Quentin-Millet and Estelle Besson (TheraVectys) for precious discussion and advices, and to Magali Tichit et Sabine Maurin for excellent technical assistance in preparing histological sections. The SARS-CoV2 variant Delta/2021/I7.2 200 was supplied by the Virus and Immunity Unit (Institut Pasteur, Paris, France) headed by Olivier Schwartz.

This work was supported by TheraVectys and Institut Pasteur.

## Author Contributions

Study concept and design: BV, JL, CG, MB, LM, PC, acquisition of data: BV, JL, AN, PA, IF, FLC, FM, KN, MB, LM, construction and production of LV and technical support: AN, FM, CB, FA, histology: FG, DH, recombinant Spike protein: CP, HM, analysis and interpretation of data: BV, JL, FA, MB, LM, PC, drafting of the manuscript: LM.

