## Supplemental Information for "An intranasal lentiviral booster broadens immune recognition of SARS-CoV-2 variants and reinforces the waning mRNA vaccine-induced immunity that it targets to lung mucosa"

Benjamin Vesin^1,£^, Jodie Lopez^1,£^, Amandine Noirat^1,£^, Pierre Authié^1,£^, Ingrid Fert^1^, Fabien Le Chevalier^1^, Fanny Moncoq^1^, Kirill Nemirov^1^, Catherine Blanc^1^, Cyril Planchais^2^, Hugo Mouquet^2^, Françoise Guinet^3^, David Hardy^4^, Christiane Gerke,^5^ François Anna^1^, Maryline Bourgine^1^, Laleh Majlessi^1,$,*,^
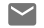
, and Pierre Charneau^1,$,*^

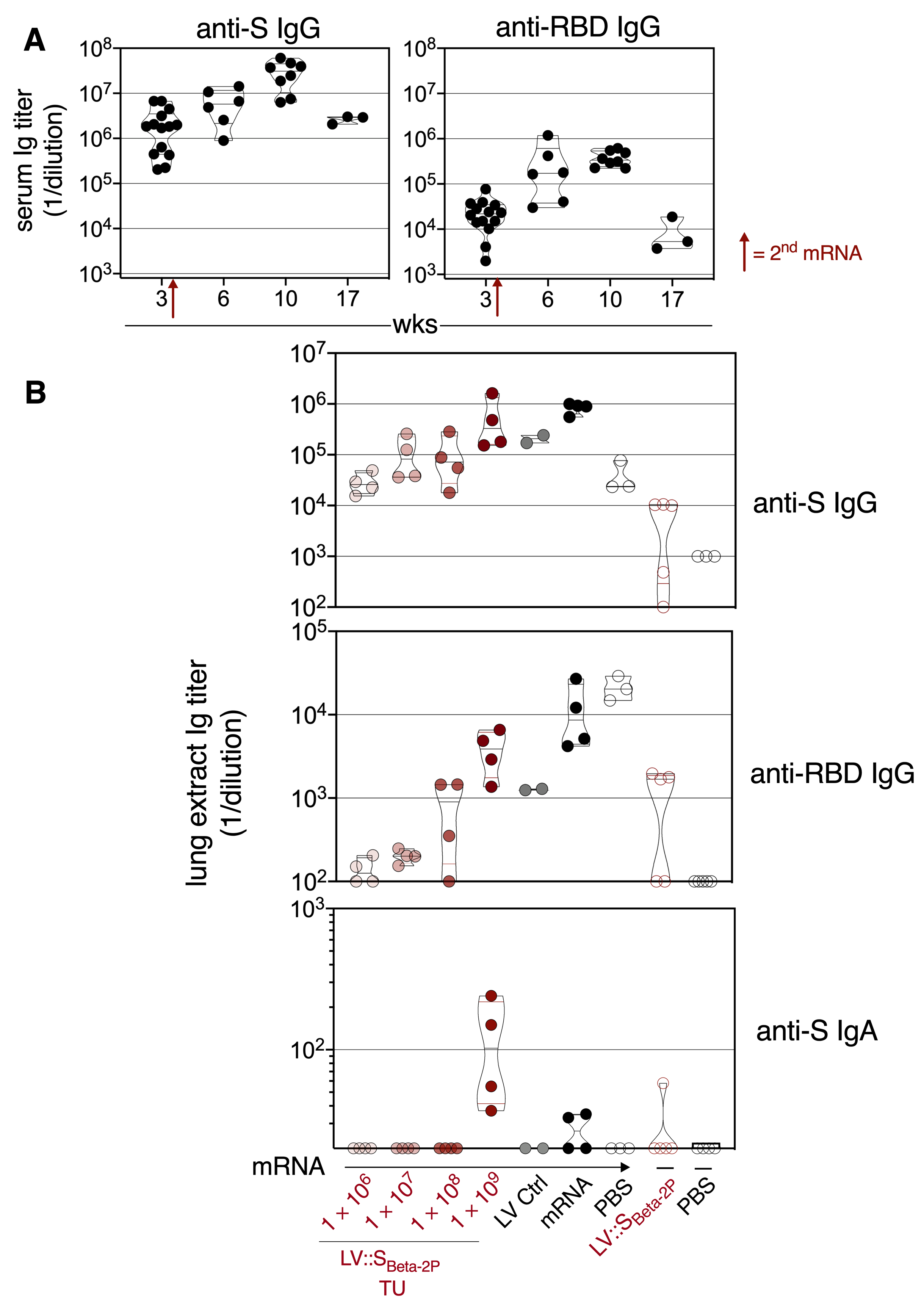

**Figure S1. Anti-S_CoV-2_ humoral responses in mRNA-vaccinated mice which were further intranasally boosted with LV::S_Beta-2P_. (A)** Follow-up of anti-S_CoV-2_ (left) and anti-RBD (right) IgG in the sera of mice initially primed and boosted i.m. with mRNA. **(B)** Anti-S_CoV-2_ IgG (top), anti-RBD IgG (middle), and anti-S_CoV-2_ IgA (bottom) in the total lung extracts of mice initially primed and boosted i.m. with mRNA and then boosted later with a third i.m. dose of mRNA or an i.n. boost of LV::S_Beta-2P_.

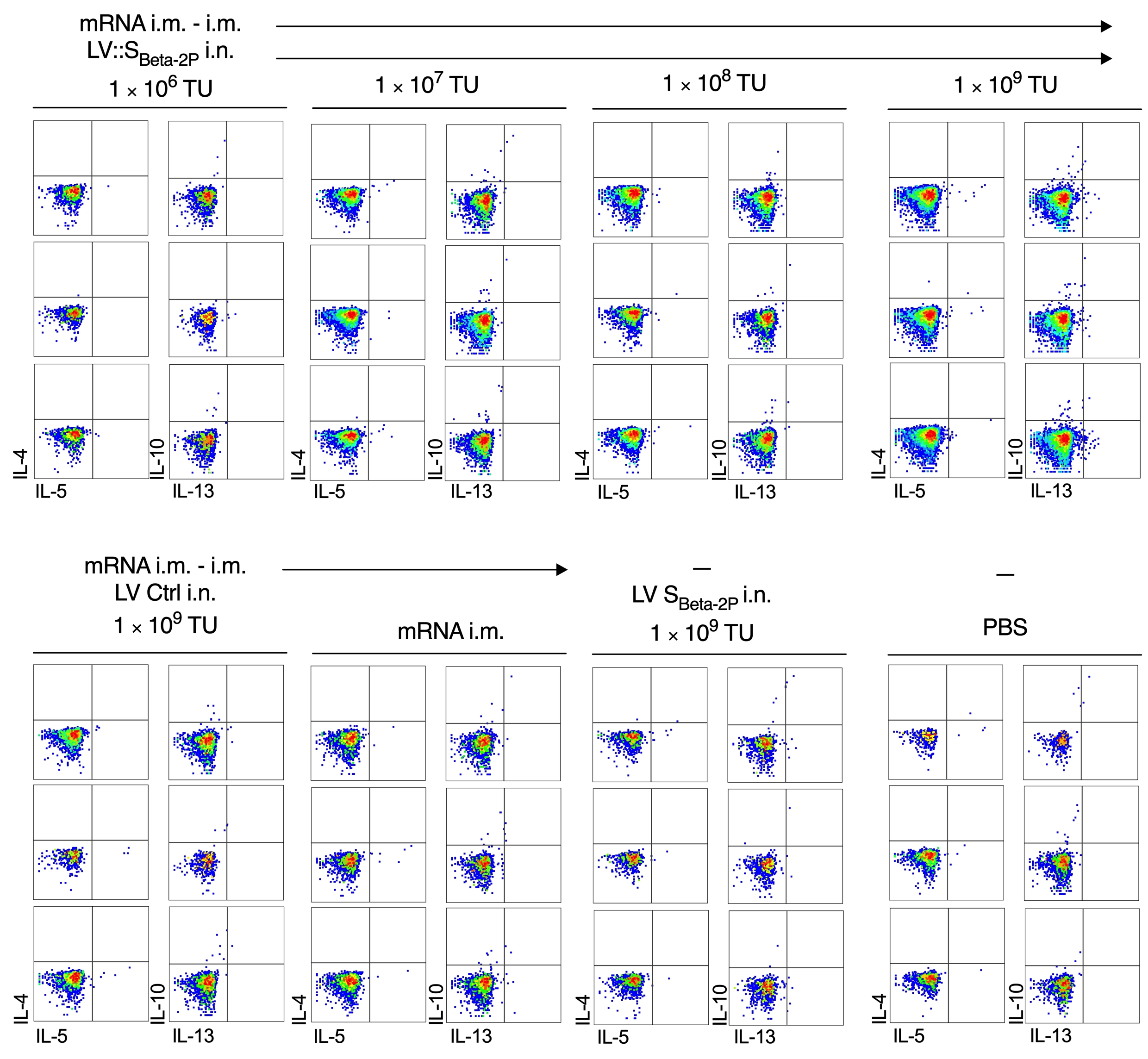
**Figure S2. Absence of mucosal CD8^+^ Tc2 responses to S_COV-2_ in mRNA-vaccinated mice which were further intranasally boosted with LV::S_Beta-2P_**. The mice are those detailed in the Figure 2. Absence of IL-4, IL-5, IL-10 and IL-13 production by lung CD8^+^ T cells after in vitro stimulation with a pool of S:256-275, S:536-550 and S:576-590 peptides, studied by ICS in parallel to the assay performed to detect IFN-γ/TNF/IL-2 (see Figure 5). Cells are gated on alive CD45^+^ CD8^+^ T cells.

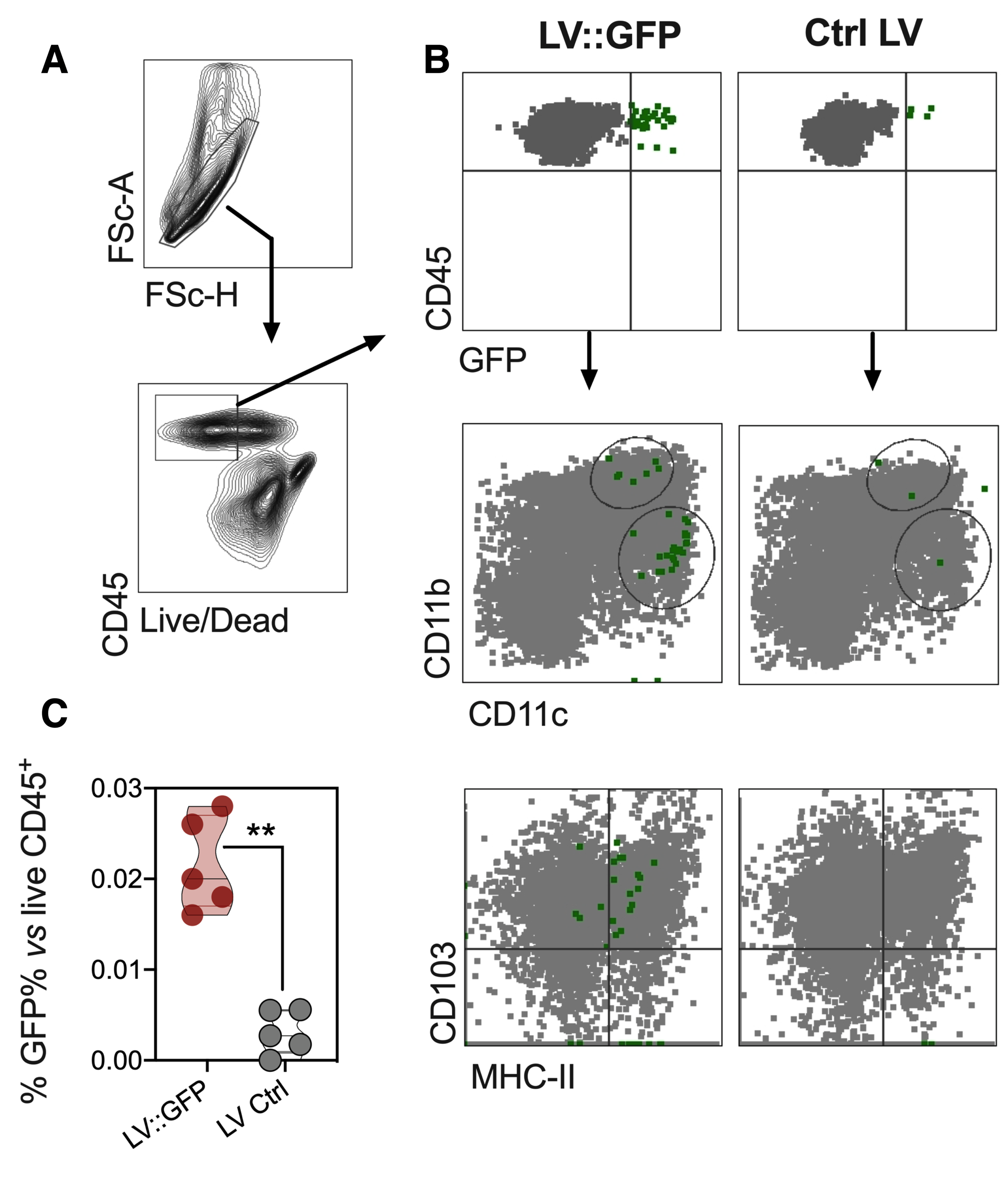

**Figure S3. Detection of cells transduced in vivo following i.n. administration**. C57BL/6 mice (*n* = 5/group) were immunized i.n. with 1 × 10^9^ TU of LV::GFP or LV::nLuc as a negative control. **(A)** Gating strategy and cytometric plots for lung immune cells from the both groups as analyzed at day 4 post-injection. **(B)** GFP^+^ cells (green) are overlaid on the total alive CD45^+^ population (gray), analyzed for the expression of other subset-specific markers. **(C)** Percentages of GFP^+^ cells versus alive CD45^+^. Statistical significance was evaluated by Mann-Whitney test (**= *p* < 0.01).

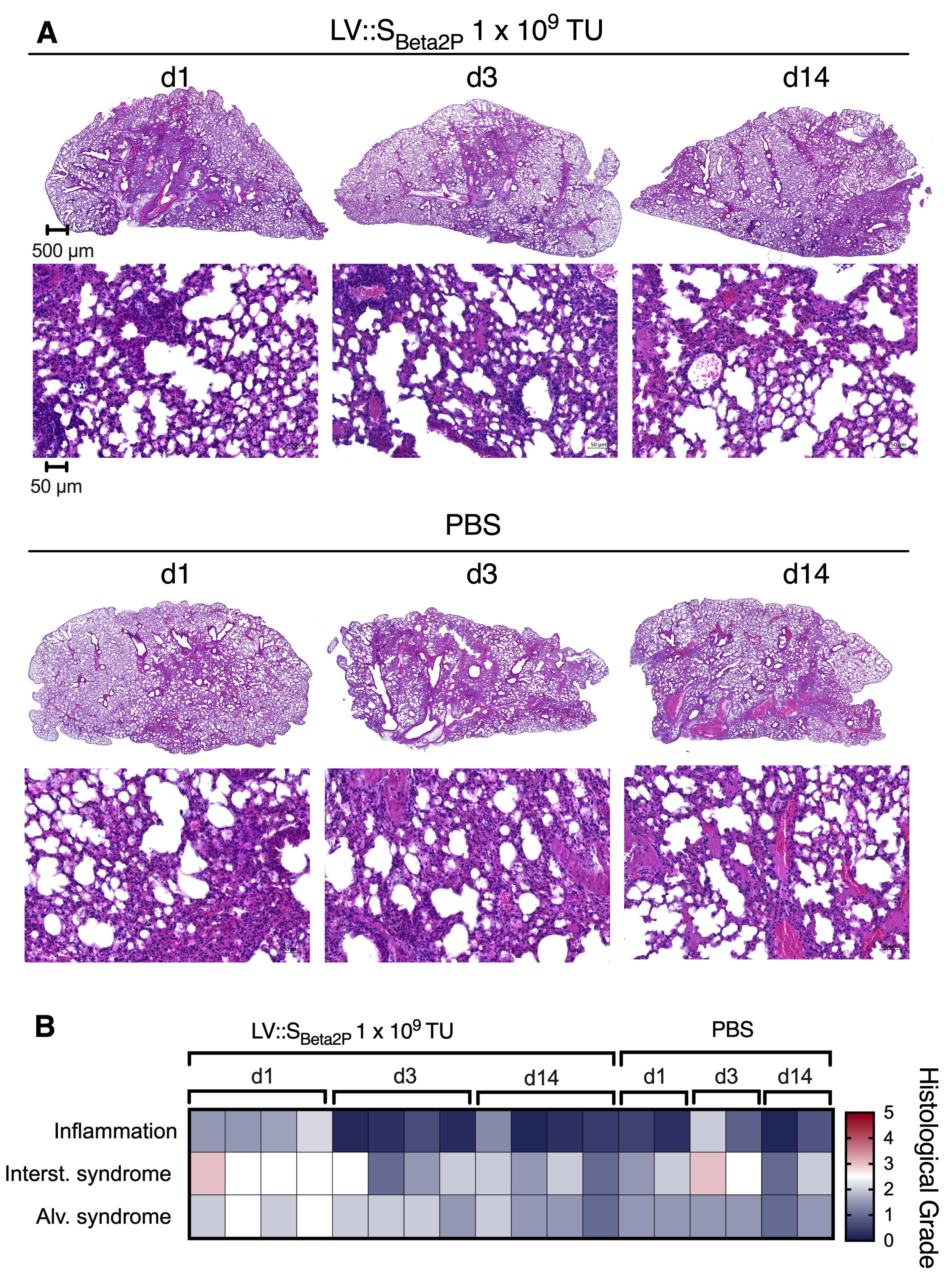
**Figure S4. Histology of the lungs after i.n. administration of LV::S_Beta-2P_.** C57BL/6 mice (*n* = 4/group) were treated i.n. with 1 × 10^9^ TU of LV::S_Beta-2P_ or PBS as a negative control. **(A)** Representative H&E whole-lung section (top, scale bar: 500 μm) and at higher magnification (bottom, scale bar: 50 μm)) at days 1, 3 and 14 post-administration. **(B)** Heatmap representing the histological scores for: (i) inflammation seriousness, (ii) interstitial (“Interst.”) syndrome, and (iii) alveolar (“Alv.”) syndrome.

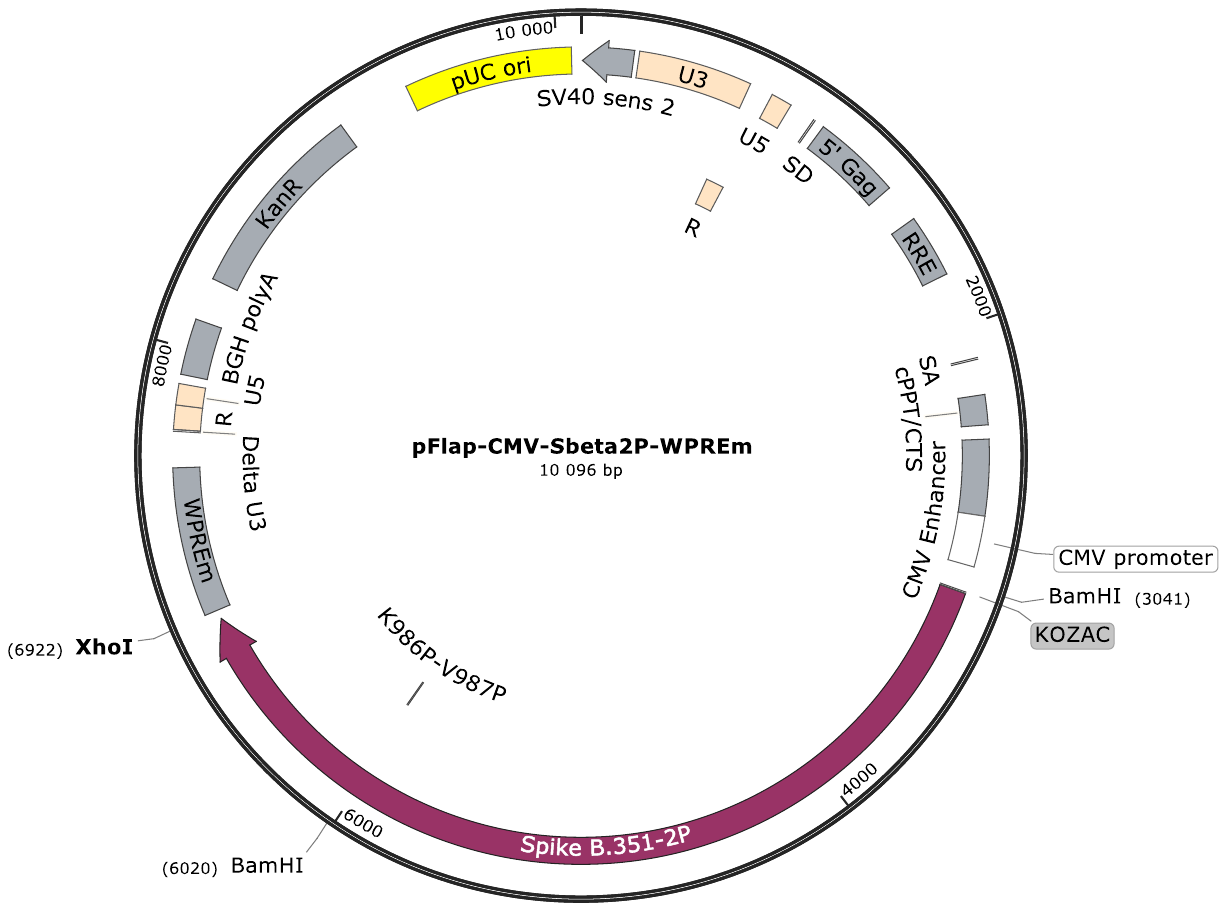

**Figure S5.** **Map of lentiviral plasmid encoding for S_Beta-2P_.** The full-length S_Beta-2P_ sequence which harbors K^986^P and V^987^P consecutive substitutions, as indicated on the map. WPREm = mutated Woodchuck Posttranscriptional Regulatory Element translational enhancer.
